# Cerebral vascular tortuosity and aneurysm formation and rupture: a novel vessel tortuosity scale

**DOI:** 10.1101/2025.06.24.661364

**Authors:** Maxon V. Knott, Naser Hamad, Pedro B. Rodrigues, Sai Sanikommu, Jennifer S. Suon, Helena Hernandez-Cuervo, Jayro Toledo, Ahmed Abdelsalam, Tiffany A. Eatz, Victor J. Del Brutto, Evan M. Luther, Sanjoy K. Bhattacharya, Joshua M. Hare, Michal Toborek, Roberto I. Vazquez-Padron, John W. Thompson, Robert M. Starke

## Abstract

Cerebral aneurysm (CA) rupture is the most common cause of nontraumatic subarachnoid hemorrhage. Recent data suggests that tortuosity is associated with aneurysm formation and rupture risk. We aimed to determine if tortuosity correlates with CA development and rupture in a mouse CA model and to develop a novel tortuosity scale to be used for *in vivo* CA studies. A highly validated, elastase-mouse CA model was used to assess cerebral vessel tortuosity with CA formation and rupture in sham and elastase groups. A 4-point ordinal scale was created to evaluate predictive capacity for vessel tortuosity level and CA formation and rupture. Nearly all sham animals (92%) had little to no vessel tortuosity on the visual scale (median, IQR: 1, [1-2]), compared to 24% in the elastase groups (2, [2-3]) (p=0.001). Sham cohorts had zero animals with highly tortuous vessels, while 3.5mU and 35mU cohorts had >35% of animals with significant visual tortuosity, p=0.003 and p<0.000, respectively. CA formation and rupture was higher in the elastase groups compared to the sham group (p=0.002). Both the visual scale and tortuosity index significantly predicted CA formation (p<0.001) and rupture (p<0.001). A novel tortuosity scale is highly predictive of CA formation and rupture *in vivo*. It may offer a new measurement to better understand vessel stress in the pathogenesis and progression of CAs.

## INTRODUCTION

Cerebral aneurysms (CA) are a focal dilation of a cerebral artery. Unruptured CAs occur in approximately 3% of the global adult population. CAs represent a critical health concern due to their potential to spontaneously rupture, resulting in a subarachnoid hemorrhage (SAH) with ensuing significant morbidity and mortality.^1^ Despite advancements in research and diagnostic strategy, the ability to predict aneurysm formation and rupture risk remains a challenge.

Structural tortuosity of cerebral vasculature has emerged as a potential factor in aneurysm formation and progression as well as in other cerebral and non-cerebral vascular diseases.^2–5^ Vessel tortuosity, defined as abnormal curvature or twisting of vessels, can result from many factors such as, genetic predisposition, hemodynamic changes, and aging.^6^ Tortuous vessels may contribute to CA development through alteration of blood flow patterns, increased wall stress, and endothelial cell dysfunction, which is associated with CA development.^7–11^

To facilitate the study of vessel tortuosity, quantitative grading scales have been developed, incorporating parameters such as vessel curvature, diameter, and branching angles.^2, 12^ These metrics enable standardization and cross-study comparisons of vascular tortuosity. Mouse models have been widely used to study CA development and progression.^13^ But few studies measure or discuss vessel tortuosity, an indicator of vessel stress.^14^ Therefore, in this study, we used a highly validated elastase induced-CA mouse model to characterize the relationship between cerebral vessel tortuosity and aneurysm formation and rupture.^15^ A novel, user-friendly visual scale was also developed to enable quantitation of vessel tortuosity.

## MATERIAL AND METHODS

All animal procedures were performed under the approval of the University of Miami Institutional Animal Care and Use Committee and our study adhered to the ARRIVE guidelines. All surgical procedures were performed under the survival plane of anesthesia with isoflurane. Confirmation of proper induction was confirmed with lack of response to noxious stimuli. Analgesia was accomplished via sustained-release buprenorphine and was administered prior to each surgical procedure.

### Mouse CA Formation

CA were induced in 8- to 10-week-old C57BL6 mice using a highly validated hypertensive-elastase aneurysm model.^16–18^ To induce hypertension, mice underwent unilateral nephrectomy followed by implantation of a deoxycorticosterone acetate (DOCA) pellet (Innovation Research of America) one week later. On the same day as DOCA pellet placement, animals were started on drinking water containing 1% NaCl. To induce aneurysm formation, mice underwent a single stereotactic injection of elastase (3.5mU or 35mU, Worthington Biochemical) into the right basal cistern at the same time as DOCA pellet placement. Sham control mice were treated as above (nephrectomy, DOCA pellet, and 1% NaCl) but received a single stereotactic injection of PBS instead of elastase. Mice (n=42) were randomly assigned to sham (n = 13), 3.5mU (n = 14), or 35mU (n = 15), elastase treatment groups. Blood pressures were measured before and at 7-, 14-, 21-, and 28-days following aneurysm induction using a tail cuff-based method (CODA Monitor Noninvasive blood pressure, Kent Scientific).

Neurological examination was performed daily to detect CA rupture. Neurological symptoms were graded as follows: (0) normal; (1) reduced eating and drinking activity demonstrated by a weight loss greater than two grams of body weight over 24 hours; (2) flexion of the torso and forelimbs upon lifting of the whole animal by tail; (3) circling to one side with a normal posture at rest; (4) leaning to one side at rest; and (5) no spontaneous activity. Mice were promptly euthanized if they developed neurological symptoms with a score of (1 - 5). All asymptomatic mice were sacrificed 28 days after aneurysm induction. Inclusion and exclusion criteria were set prior to study initiation. Exclusion criteria included mice who expired prior to euthanasia or those with failed transcardial perfusion that did not allow visualization of cerebral vasculature. All mice euthanized based on the prior mentioned guidelines, with successful transcardial perfusions were included in analysis.

To determine CA formation and rupture rates, the mice underwent transcardial perfusion with phosphate-buffered saline (PBS), followed by a 20% gelatin and 0.1% methylene blue solution to visualize the cerebral arteries. Both whole brain and magnified images of the cerebral vessels were obtained and the images analyzed for CA formation, rupture, and vessel tortuosity by two blinded observers. An aneurysm was defined as an outward bulging of the vessel wall, 1.5 times the diameter of the parent artery

### Vessel Tortuosity Index Measurements

Vessel tortuosity of the anterior communicating artery (ACA), middle cerebral artery (MCA), and basilar artery (BA) were determined by calculating the Tortuosity Index (TI) for each vessel. The TI is defined as the ratio of the actual vessel length between two points divided by the length of a straight line between the same two points, TI = Distance actual / Distance straight. ImageJ software from NIH was used to calculate vessel distances on magnified images of the target vessel between the vessel origin and terminus. Each vessel was measured three times by 2 independent observers, blinded to treatment, a third observer was utilized to ensure inter-rate reliability. The average vessel measurement was determined and used for the study analysis.

### Qualitative Vessel Tortuosity Measurement

Qualitative vessel tortuosity measurements were initially determined by the degrees the vessel deviated from its intended path. A line connecting the vessel before and after its tortuous segment was used to visualize the angle of deviation. The degrees to which the vessel deviated was the sum of the total measurements. Measurements were then placed into four categories: 0-44°, 45-89°, 90-120°, and 121-180°.

Large and small vessel tortuosity was then further simplified by visualization of curvature. Frank tortuosity was identified to be most clear when the vessel deviated from its path >45°. Large vessels included the MCA, ACA, and BA; clearly visible tortuosity in these vessels was termed “Large Turns” in our visual scale. Likewise, all non-large vessels were considered small vessels, and clearly visible tortuosity in these vessels was termed “Small Turns.” A vessel’s final visual scale score was determined by the summation of large turns and small turns, which then could be correlated with its respective categories.

### Statistics

All data are expressed as mean ± S.D. Statistical analysis between two groups was performed using the unpaired Student’s t-test. Statistical analysis between more than two groups was performed using a one-way ANOVA with Dunnett’s multiple comparison post hoc test. A p-value less than 0.05 was considered statistically significant. Pearson’s correlation coefficient was then applied to analyze the relationship between TI and aneurysm formation and rupture.

## RESULTS

### Group Descriptions and CA Model Confirmation

The presented study consists of 46 total animals, with 13 (28%) in the sham, 14 (30%), in the 3.5mU, and 19 (41%) in the 35mU cohorts. Analysis was performed using the entire cohort unless otherwise stated in figure legends. CA formation was carried out as previously described using a single stereotactic elastase injection (3.5 and 35mU) to induce aneurysm formation. CA formation and rupture rates were determined 28 days later or upon development of neurological score of 1-5. Cerebral aneurysms were not observed in sham-treated animals but were formed and spontaneously ruptured in a dose dependent fashion. In both the 3.5mU and 35mU elastase injection groups, 13%, and 27% of animals demonstrated unruptured aneurysm formation while 20% and 50% experienced ruptured CA (Figure 1A and B), respectively.

**Figure 1:**
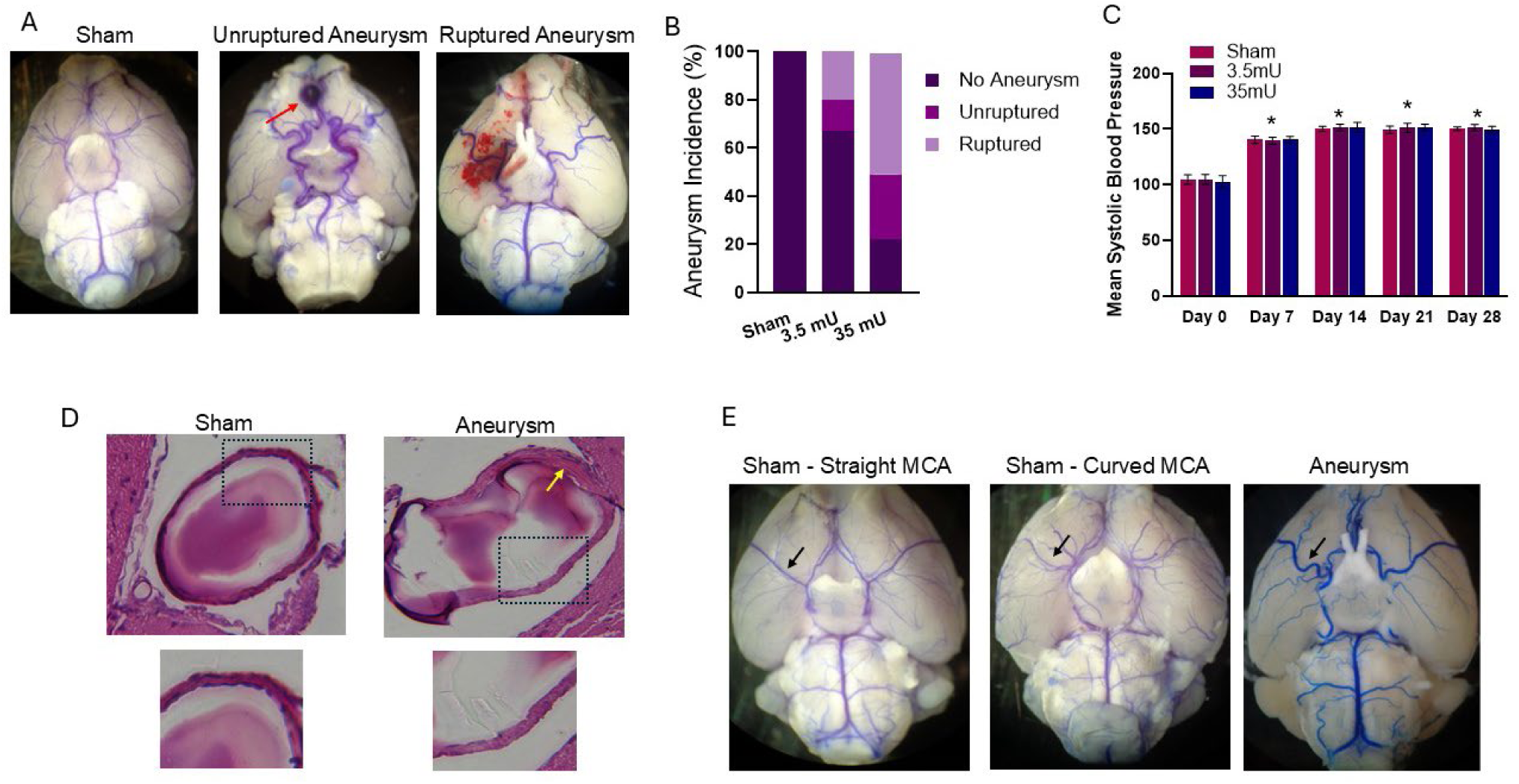
CA formation and rupture rates. (A) Light microscopic representative images of bromophenol blue infused cerebral blood vessels in sham, unruptured aneurysm, and ruptured aneurysm mice. (B) Quantitation of CA formation in sham (0%), 3.5mU (13%), and 35mU (27%) groups and rupture in sham (0%), 3.5mU (20%), and 35mU (50%). (C) Hypertension was equally increased in Sham and CA cohorts (n = 7). (D) H&E staining of normal cerebral blood vessel in sham operated mice showing a thin continuous layer of endothelial cells but discontinues in the areas of the thin vessel walls in the elastase treated mice. (E) Representative images taken via light microscopy of cerebral blood vessels show normal and gently curved vessels in sham-operated mice. Additionally, a representative image in the elastase-treated mice group shows severe tortuosity of the vessels. Black arrows point to variations in the right MCA. \**P*<0.05 vs day 0 using Kruskal–Wallis analysis with Wilcoxon rank-sum post hoc test.

To confirm hypertension was induced in all mouse cohorts, we used a non-invasive tail cuff blood pressure measurement to monitor hypertension 7, 14, 21 and 28 days following DOCA pellet placement and starting 1% sodium chloride drinking water. Blood pressure was measured prior (Day 0) to hypertension induction to establish baseline blood pressure measurements. As shown in Figure 1D, blood pressure was significantly elevated at 7 days and remained elevated for the duration of the experiment. There was no difference found in blood pressure between sham and elastase treated cohorts at any of the times measured.

As previously reported, H&E assessment of cerebral vessels revealed distinct changes in CA vessel wall architecture (Figure 1E). Vessels from sham cohorts displayed a continuous endothelial layer and two to three layers of smooth muscle cells, whereas in CA there was a discontinuous layer of endothelial cells, indicating a disruption in endothelial integrity and scattered and stenotic vascular smooth muscle cells. These findings are consistent with prior studies that reported similar discontinuities in the vascular endothelial layer following elastase treatment and are consistent with that observed in human CA.^19^

### Quantitative Tortuosity Assessment and CA formation/rupture

In addition to CA formation and rupture, we found that 3.5mU and 35mU elastase treatment induced a varying degree of cerebral vessel tortuosity when compared to sham control cohorts (Figure 1F). We also observed variability in the right angulation MCA of both sham and elastase cohorts, ranging from straight to having a mild curvature. While in CA cohorts more acute changes in vessel directions were observed.

Therefore, to better understand the association of vessel tortuosity in CA pathology, we determined the incidence of at least one acute vessel change greater than 45 degrees in the ACA, MCA, and BA. Since elastase is injected into the right basal cistern, for these initial studies we limited our analysis to only the right ACA and MCA. As shown in Figure 2A-C, there was only one sham cohort with MCA vessel tortuosity greater than 45 degrees. No other vessel tortuosity was observed in sham cohorts. In contrast, there was a significant increase in vessel tortuosity in animals which underwent CA surgery with tortuosity frequency being observed in the MCA at 50% and 55%, in the ACA at 36% and 35%, and in the BA at 38% and 20% for 3.5mU and 35mU cohorts, respectively. To better characterize the severity of tortuosity, we then determined the frequency of angles observed between 0-44, 45-89, 90-120, and 121-180 degrees in the right MCA and ACA, and BA. Representative angles are depicted in Figure 2A. As shown in Figure 2E-G, the majority of observed angle changes greater than 45 degrees were between 121-180 degree in all three vessels analyzed.

**Figure 2:**
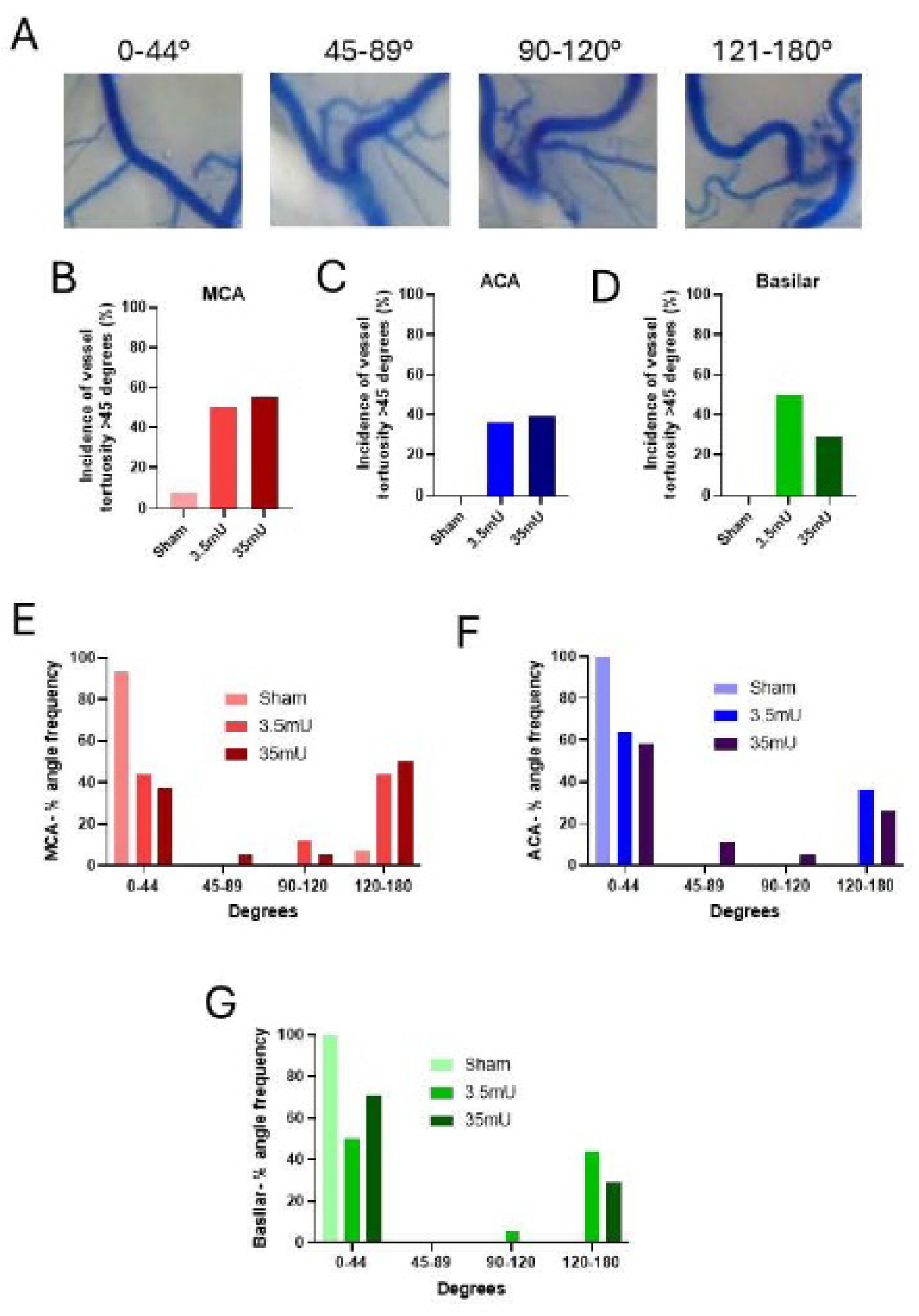
Incidence of vessel tortuosity and distribution of angle variance between groups for major vessels. (A) Representative vessel angle grouping. (B) Incidence of right MCA vessel tortuosity (>45 degrees) was highest in the 35mU group (55%), followed by the 3.5mU group (50%), and each elastase group had significantly higher incidence than the sham group (7.7%). (C) Incidence of ACA vessel tortuosity (>45 degrees) was highest in the 3.5mU group (36%, 5/14), followed by the 35mU group (35%, 6/17), and each elastase group had significantly higher incidence than the sham group (0%, 0/10). (D) Incidence of basilar vessel tortuosity (>45 degrees) was highest in the 3.5mU group (38%, 5/13), followed by the 35mU group (20%, 3/15), and each elastase group had significantly higher incidence than the sham group (0%). (E) The sham cohort had the greatest incidence of MCA vessel angles in the 0– 44-degree category (93%), whereas the 3.5mU and 35mU had the greatest incidence of MCA vessels measuring 120-180 degrees (42% and 50%, respectively). (F) The sham cohort had the greatest incidence of ACA vessel angles in the 0–44-degree category (100%), whereas the 3.5mU and 35mU had the greatest incidence of ACA vessels measuring 120-180 degrees (36% and 28%, respectively (G) The sham cohort had the greatest incidence of basilar vessel angles in the 0–44-degree category (100%), whereas the 3.5mU and 35mU had the greatest incidence of basilar vessels measuring 120-180 degrees (50% and 31%, respectively).

Next, we wanted to determine if vessel tortuosity correlated with CA formation and rupture. Therefore, we measured the tortuosity index (TI), which is defined as the actual vessel length between points A and B divided by a straight line between the same points (Figure 3A and B).^20^ As shown in Figure 3C, the right MCA TI was significantly increased approximately 20% with both 3.5mU and 35mU elastase treatment compared to sham cohorts. There was no difference in MCA TI between 3.5mU and 35mU elastase treated cohorts. Despite observed vessel tortuosity in the right ACA and BA (Figure 2), the TI was not significantly increased over sham cohorts by either elastase concentrations (Figure 3C-E). Next, we determined the correlation between right MCA and ACA, and BA TI to the incidence of CA formation and rupture using Pearson’s correlation coefficient. As shown in Figure 4A-F, we found that there was no significant correlation between TI and CA formation and rupture in either the ACA or BA, but we did find a significant correlation between MCA TI and CA formation (p = 0.0369) and rupture (p = 0.0372).

**Figure 3:**
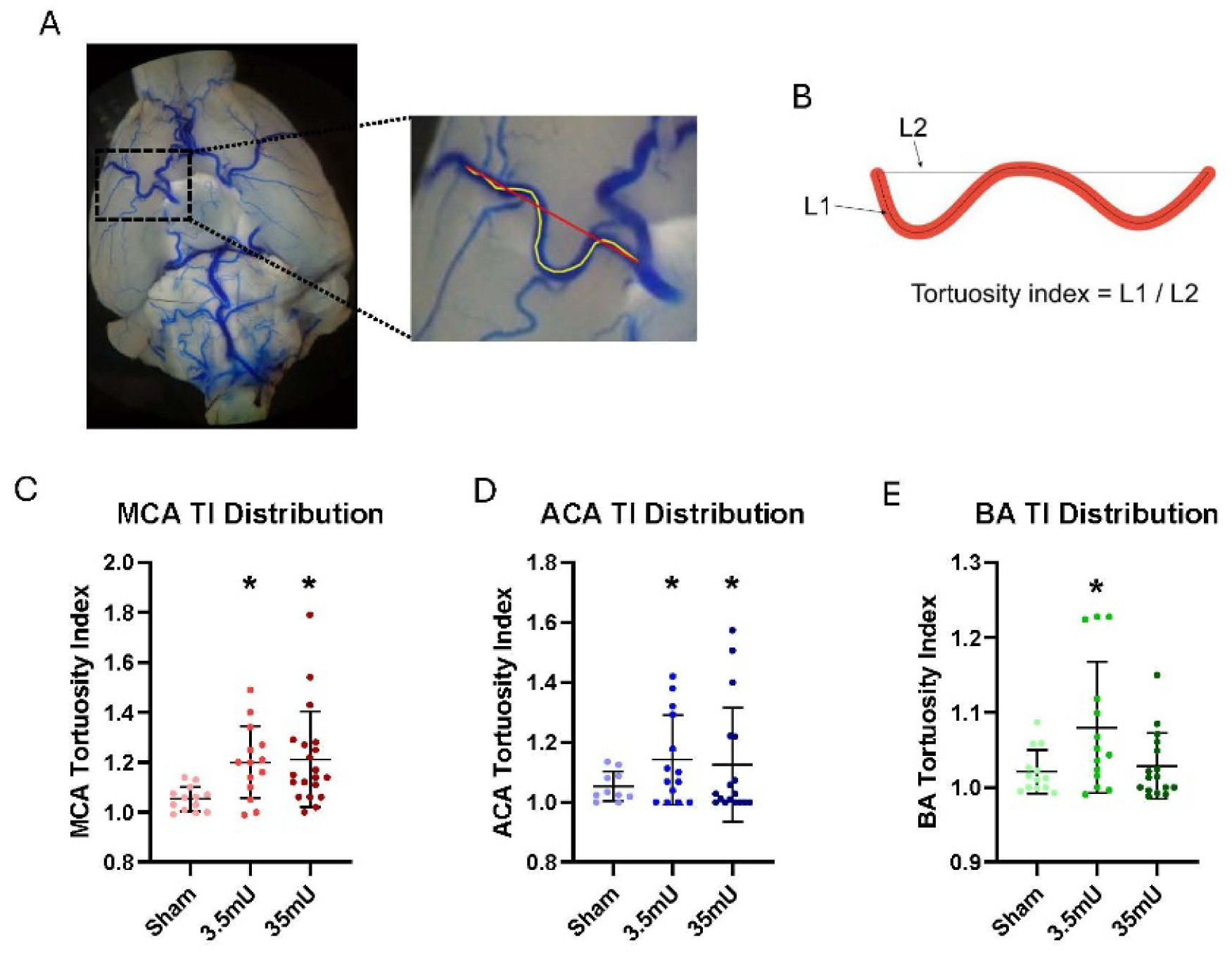
MCA TI distributions across groups. (A) The tortuosity index was measured from the take-off points of the MCA to the end point before tissue curvature. (B) TI was calculated by L1/L2 with L1 being a direct vessel tracing and L2 being a straight line from vessel take-off to end-point. (C) An unpaired t-test with Welch’s correction found that the MCA TI was significantly higher in the 3.5mU group (M=1.200) when compared to the sham group (M=1.053) (Difference between means +/− SD: 0.15 +/− 0.04, p=0.0025. Similarly, the MCA TI was significantly higher in the 35mU group (M=1.211) when compared to the sham group (M=1.053) (0.16 +/− 0.05, p=0.0019). There was no difference between 3.5mU and 35mU MCA TI. ACA and basilar TI are shown in D and E.

**Figure 4:**
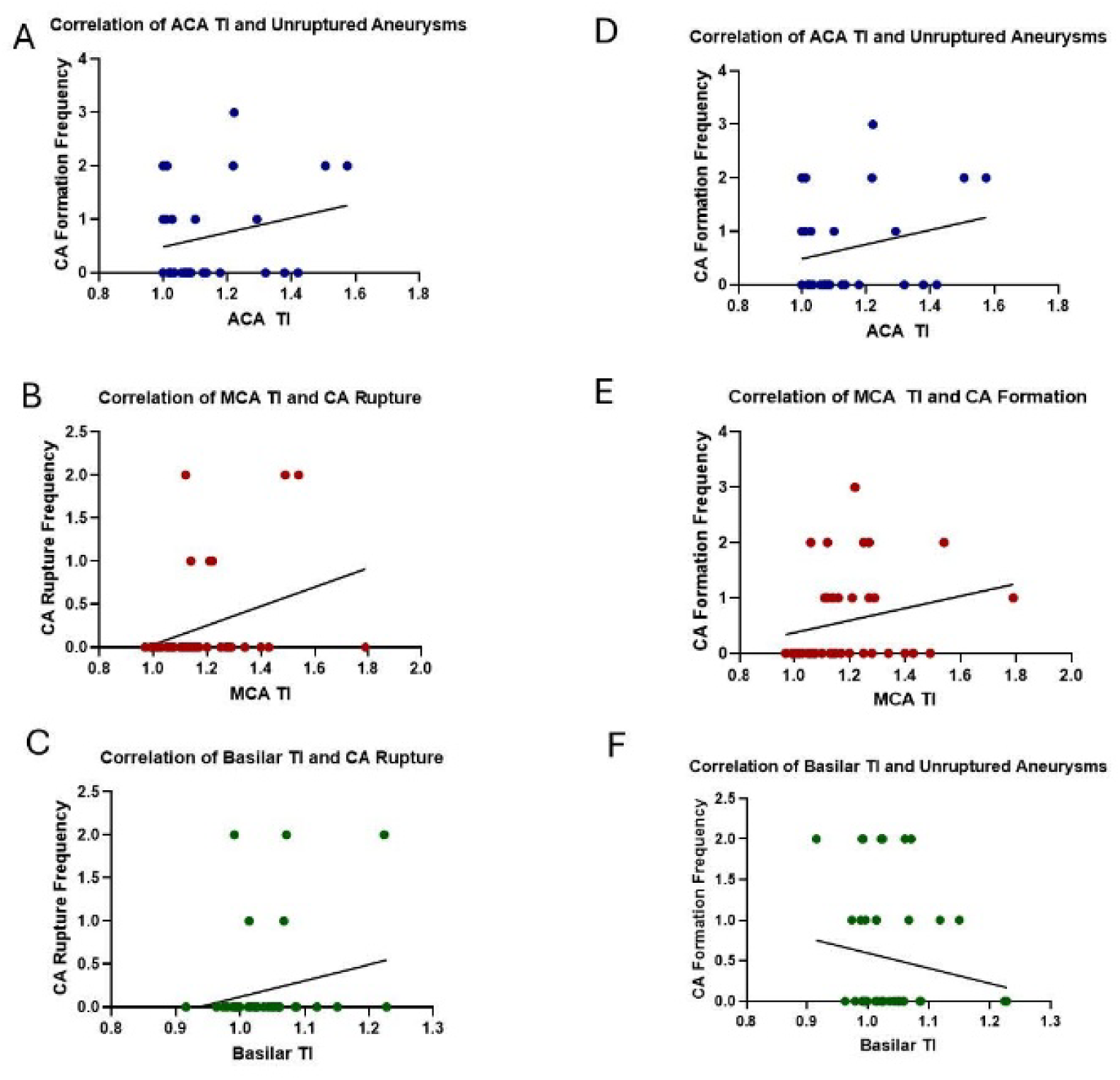
Pearson’s correlation coefficient analysis of large vessel TI and ruptured CA in A: ACA, B: MCA, and C: BA; correlation of large vessel TI and unruptured CA in D: ACA, E: MCA, and F: BA. No significant correlations resulted between TI and CA formation and rupture in either ACA or BA; but significant correlation was calculated between MCA TI and CA formation (p = 0.0369, R = 0.3052) and rupture (p = 0.0372, R = 0.3049).

### Qualitative Tortuosity Measurement

Since elastase is injected into the right basal cistern, the majority of CA formation and rupture are observed on the right side of the brain. However, CA formation and rupture are also found on the left side of the Circle of Willis as well, just at a lower frequency. Therefore, we determined the incidence of bilateral tortuosity and found that 71% of 3.5mU and 36% of 35mU elastase treated cohorts had bilateral MCA tortuosity (Figure 5A and B). Similarly, the right MCA TI was significantly higher in animals with bilateral vessel tortuosity compared to those without (p<0.0001) (Figure 5C).

**Figure 5:**
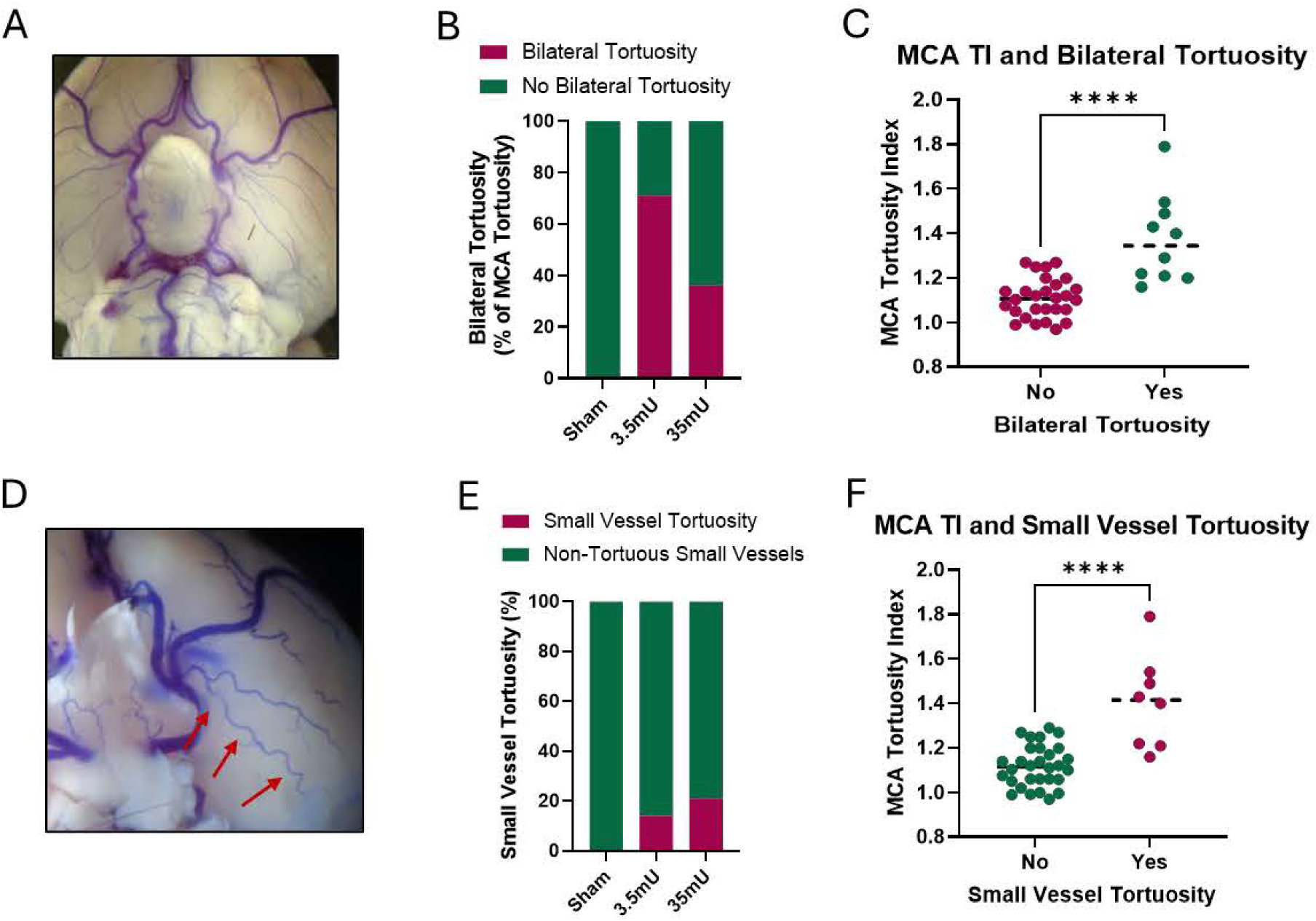
Bilateral and Small vessel tortuosity. (A) Bilateral tortuosity was defined as large vessels having tortuous angles in each hemisphere. (B) Bilateral tortuosity was present in 26% of the 35mU, 42% (5/12) of the 3.5mU, and 0% of the sham animals. (C) A Mann-Whitney t-test found that the MCA TI was significantly higher in animals with bilateral vessel tortuosity and those without (p<0.0001). (D) Small vessel tortuosity, as demonstrated by the red arrows, was defined in non-large cerebral vessels with multiple turns. (E) The 35mU group had the greatest incidence of small vessel tortuosity (21%), as compared to the 3.5mU group (14%), and the sham group (0%). (F) A Wilcoxon test found that the MCA TI was significantly higher in animals with small vessel tortuosity and those without (p=0.0078).

It was also observed that small vessels displayed notable tortuosity (Figure 5D) in elastase treated cohorts. We found no instance of small vessel tortuosity in sham animals but 15% and 20% of elastase treated cohorts exhibited small vessels tortuosity in the low and high dose elastase groups, respectively (Figure 5E). The right MCA TI was significantly higher in animals with small vessel tortuosity compared to those without (p<0.0001) (Figure 5F).

### Tortuosity and CA formation/rupture

Our data demonstrates that the incidence of vessel tortuosity is increased in CA animals and that MCA tortuosity correlates with CA formation and rupture. Therefore, to identify a reliable indicator of vessel stress associated with CA formation, we developed a tortuosity scale ranging from 1-non-tortuous to 4-major vessel tortuosity (Figure 6A). This scale was designed to visually assess the level of tortuosity based upon the number of large and small vessel turns without the need of vessel measurements. To validate the effectiveness of this scale, we conducted several tests within our experimental model. As shown in Figure 6B, similar to the TI in Figure 3, the visual tortuosity score was significantly increased in 3.5mU and 35mU elastase treated cohorts when compared to sham controls. The tortuosity scale was also demonstrated to predict the incidence of CA formation and rupture (Figure 6 C and D). These results demonstrate that our visual tortuosity scale is reliable and replicates the degree of vessel tortuosity determined by more analytical methods.

**Figure 6:**
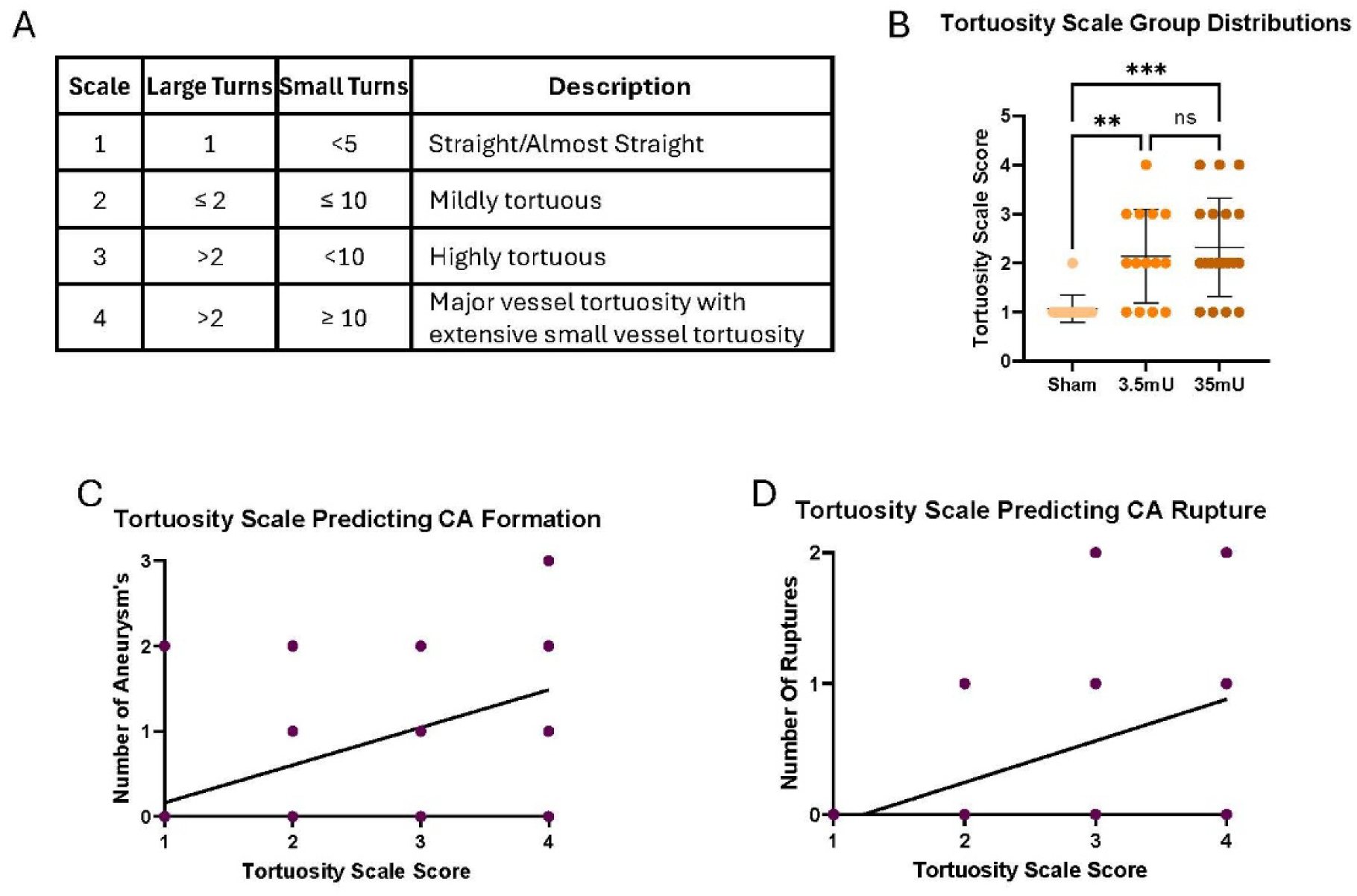
Tortuosity scale predicts the incidence of CA formation and rupture. (A) Tortuosity scale consisting of the number of large and small vessel turns with an ordinal range of 1-4. (B) Visual scale differed significantly between the three groups via one-way Anova [F (2,43) = 9.028, r^2^=0.30, p=0.0005]. Multiple comparisons showed that the sham group had significantly lower values on the visual scale than the 3.5mU group (Mean difference, 95% CI: −1.07 [−1.9, −0.3], p=0.0058) and the 35mU group (−1.2 [−2.0, −0.5], p=0.0006). The mean visual scale scores did not differ between the 3.5mU and 35mU groups (p=0.831). (C-D) The institutional visual scale and tortuosity index significantly predicted CA formation (p<0.001) and rupture (p<0.000).

**Figure 7:**
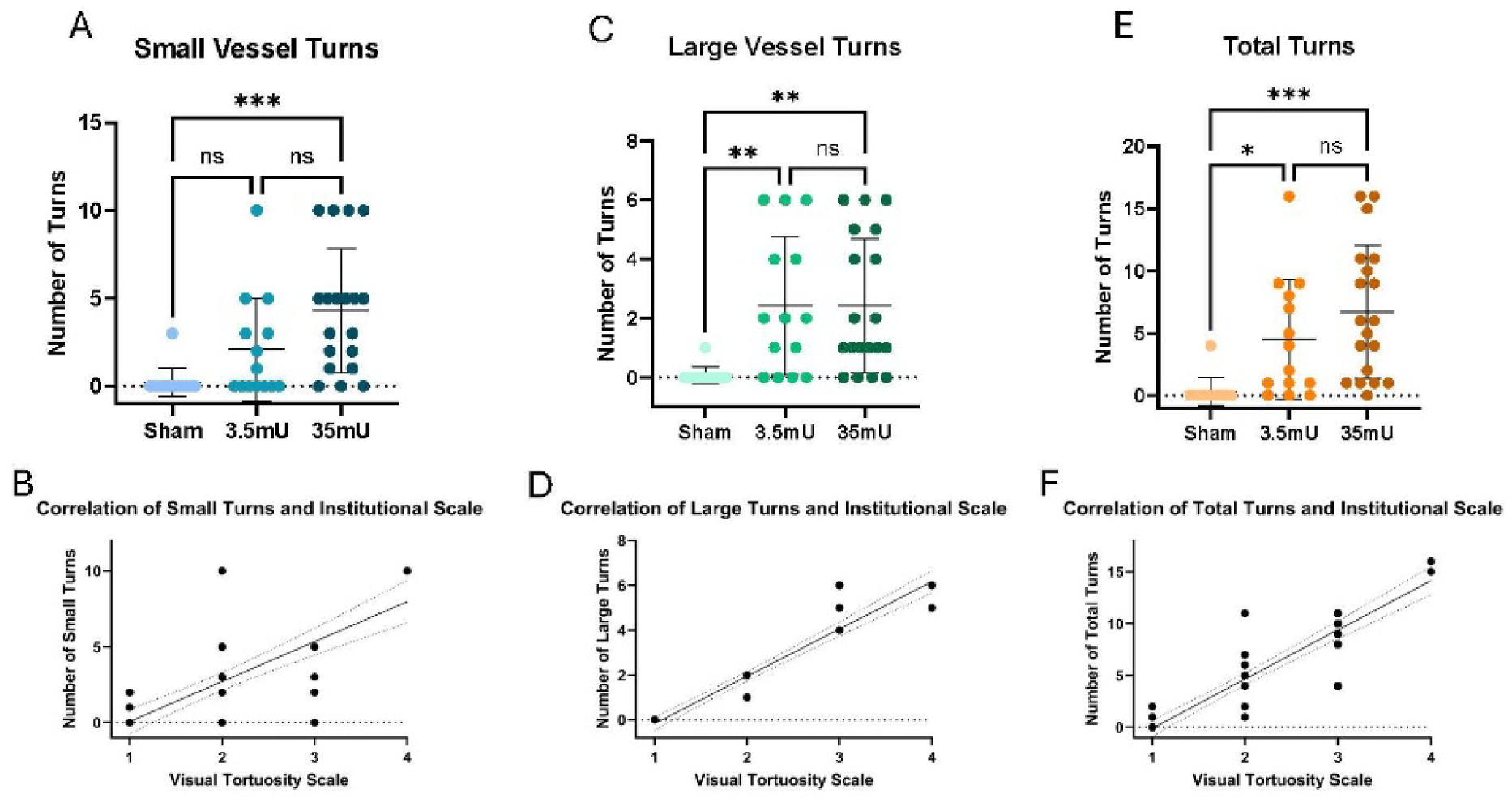
Group distributions of the tortuosity scale and its components. (A) Small vessel turns differed significantly between the three groups via one-way Anova [F (2,43) = 8.23, r^2^ =0.28, p=0.0009]. Multiple comparisons showed that the sham group did not have significantly lower small turns than the 3.5mU group (−1.84 [−4.5, 0.8], p=0.222), but did have less than the 35mU group (−4.085 [−6.6, −1.6], p=0.0007). The number of small vessel turns trended towards being significantly less in the 3.5mU group than the 35mU group (p=0.07) (B) The number of small turns significantly correlated with higher scores on our visual scale (r=0.74, p<0.000). (C) Large vessel turns differed significantly between the three groups via one-way Anova [F (2,43) = 6.70, r^2^ =0.24, p=0.0029]. Multiple comparisons showed that the sham group had significantly lower large turns than the 3.5mU group (−2.35 [−4.2, −0.5], p=0.009) and the 35mU group (−2.34 [−4.1, - 0.6], p=0.005). However, the number of large turns did not differ between the 3.5mU and 35mU groups (p>0.99). (D) The number of large turns significantly correlated with higher scores on our visual scale (r=0.98, p<0.000). (E) Total turns differed significantly between the three groups via one-way Anova [F (2,43) = 8.34, r^2^ =0.28, p=0.0009. Multiple comparisons showed that the sham group had significantly lower total turns than the 3.5mU group (−4.19 [−8.3, −0.1], p=0.0439) and the 35mU group (−6.43 [−10.3, −2.6], p=0.0006). However, the number of total turns did not differ between the 3.5mU and 35mU groups (p=0.33). (F) The number of total turns significantly correlated with higher scores on our visual scale (r=0.92, p<0.000).

After analyzing the major large vessels and small vessels in the brain, we quantified the number of turns in each vessel group to identify any significant differences. We then combined the turn counts from both vessel types to test if there were significant correlations within each elastase-treated group. Statistical analysis revealed significant differences in small, large, and total turns among the groups. Total turns were strongly correlated with higher scores on our visual scale (r = 0.92, p < 0.0001). These findings indicate that our visual scale is a reliable and easily implemented method for evaluating vessel tortuosity, effectively identifying significant patterns of vessel tortuosity associated with CA formation in preclinical models and providing a viable alternative to more expensive neuroimaging and more laborious image analysis techniques.

## DISCUSSION

The results of this study demonstrate that vessel tortuosity is increased in a commonly used mouse elastase CA model.^16–18^ The most common vessel tortuosity observed ranged between 120-180 degrees. Our findings also demonstrate that our visual scale (ranging from 1-4) is a reproducible, reliable, and easy method to evaluate vessel tortuosity. It effectively identified significant patterns of vessel tortuosity associated with CA formation and rupture and provides a viable alternative to more expensive neuroimaging and laborious image analysis techniques.

Consistent with previous findings, we found that CA formation and rupture rates were increased with increasing elastase concentration. Surprisingly though, vessel tortuosity, as measured by the incidence, tortuosity index, and tortuosity scale were independent of the elastase concentration used. Tortuosity was observed only in elastase treated and not sham cohorts suggesting an elastase dependent effect and not hypertension alone, which sham animals were also exposed to as well. These findings are distinct from the role of hypertension alone in CA formation which has been well documented, as the mouse aneurysm model has shown to require the synergistic effects of elastase and hypertension.^21, 22^

Using transcranial doppler ultrasound, Lebas et al. also found vessel tortuosity in the Circle of Willis in the Hosaka at al and Nuki et al. mouse CA models.^23^ The Nuki model is similar to that used in this study while the Hosaka model increases hemodynamic stress and vessel weaking by the addition of right renal artery and common carotid artery ligation and β-aminopropionitrile administration in the drinking water.^18, 24^ The incidence of vessel tortuosity in the Nuki model, was similar to that reported here. However, vessel tortuosity was greatly enhanced in the Hosaka model with 100% of animal have vessel tortuosity in the Circle of Willis by days 6 following elastase treatment. Yet the methods which the Hosaka model induces HTN differ from ours as they ligated both the right renal artery (without performing a nephrectomy) and left common carotid artery.

Our data demonstrates that approximately 40% of all elastase treated cohorts displayed tortuosity greater than 45 degrees in the MCA, ACA, and BA. Tortuosity was primarily found on the right side of the Circle of Willis, correlating with elastase injection into the right basal cistern. In contrast, the TI which measures the relative change in vessel length caused by tortuosity was only significantly increased in the MCA but was not dependent upon elastase concentration. In contrast, our visual tortuosity grading scale allowed for an accurate and quantitative representation of the overall change in vessel tortuosity and was sensitive enough to correlate with CA formation and rupture.

Bilateral and small vessel tortuosity was visualized in a number of elastase treated mice. Furthermore, the MCA TI was significantly higher in mice with bilateral tortuosity and in those with small vessel tortuosity. These findings suggest that animals with greater large vessel tortuosity often have contaminant tortuosity throughout both hemispheres and into small vessels. Studies have shown a relationship between small vessel disease and internal carotid artery tortuosity in clinical patients that underwent neuroimaging.^25^ Small vessel disease progression has been implicated as an underlying factor in many vascular disorders.^26^ This influence may suggest that changes in small vessels are reflected in larger vessels, potentially modulating overall vascular dynamics and disease risk. This relationship could aid in identifying patterns of patients at risk for aneurysm complications or rupture.

As part of this study, we developed a novel visual scale with four possible ordinal values ranging from 1 to 4 with a score of 1 representing little to no tortuosity while a score of 4 representing major vessel tortuosity. The number of large vessel turns and small vessel turns together determine the scales’ value in a given case. The visual scale was not only higher in the elastase groups, but also significantly predicted CA formation and rupture. In the clinical setting, cerebral vasculature tortuosity and its extracranial influencing vessels can be analyzed via advanced methodologies, such as transcranial Doppler, carotid artery Doppler ultrasound, magnetic resonance angiography (MRA), computed tomographic angiography (CTA), and digital subtraction angiography (DSA). These modalities are costly and may prove challenging to execute, possibly involving iodinated or gadolinium contrast and potential adverse sequalae or invasive endovascular diagnostic procedures. Yet, these studies can provide insight into vessel diameter, measurements, flow velocities, and serve as a basis for further analysis of which a tortuosity index can be calculated. This clinical TI is arrived at by dividing the length of a given vessel by a straight-line distance between two points on the same vessel, as was done in our mice-model. More advanced reconstructions and techniques can even track and measure the curvature at various points of the vessel, generating a three-dimensional tortuosity map. A visual scale could provide investigators with both quantitative and qualitative benefits to incorporate vessel tortuosity tiering into their clinical evaluations. This inclusion could lead to a more comprehensive understanding of a patient’s vascular abnormalities, specifically aneurysmal in nature, and their potential consequential progression and rupture. Therefore, research into transitioning this visual scale from animal to human models may be worth further exploration.

## CONCLUSION

To our knowledge, this study is the first to demonstrate a significant correlation between a quantitative visual tortuosity scale with CA formation and rupture in an elastase mouse model. This novel tortuosity scale is highly predictive of CA formation and rupture in vivo and may offer a novel measurement to better understand vessel stress in the natural history of CAs.

## Acknowledgments

None

## Author Contributions

Maxon Knott, investigation, writing- original draft; Naser Hamad investigation, writing- original draft; Pedro B. Rodrigues investigation, writing- original draft; Sai Sanikommu investigation, writing- original draft; Jennifer S. Suon Writing - Review and editing; Helena Hernandez-Cuervo Review and editing ; Jayro Toledo Formal analysis; Ahmed Abdelsalam Formal analysis; Tiffany A. Eatz, investigation, writing; Victor J. Del Brutto Writing -Review and editing; Evan M. Luther Writing -Review and editing; Sanjoy K. Bhattacharya Writing -Review and editing; Josh M. Hare Writing - Review and editing; Michal Toborek Writing -Review and editing; Roberto I. Vazquez-Padron Writing -Review and editing; John W. Thompson Conceptualization, formal analysis, Writing original draft; and Robert M. Starke Conceptualization, Writing- Review and editing

## Data Availability

Data is available upon request.

